# Accurate prediction of site- and amino-acid substitution rates with a mutation-selection model

**DOI:** 10.1101/2024.03.02.583099

**Authors:** Ingemar André

**Affiliations:** Biochemistry and Structural Biology, Lund University, PO BOX 124, Lund, Sweden

**Keywords:** Amino acid site-rates, substitution matrices, mutation-selection model

## Abstract

The pattern of substitutions at sites in proteins provides invaluable information about their biophysical and functional importance and what selection pressures are acting at individual sites. Amino acid site rates are typically estimated using phenomenological models in which the sequence variability is described by rate factors that scale the overall substitution rate in a protein to sites. In this study, we demonstrate that site rates can be calculated accurately from amino acid sequences using a mutation-selection model in combination with a simple nucleotide substitution model. The method performs better than the standard phylogenetic approach on sequences generated by structure-based evolutionary dynamics simulations, robustly estimates rates for shallow multiple sequence alignments, and can be rapidly calculated also on larger sequence alignments. On natural sequences, site rates from the mutation-selection model are strongly correlated to rates calculated with the empirical Bayes methods. The model provides a link between amino acid substitution rates and equilibrium frequency distributions at sites in proteins. We show how an ensemble of equilibrium frequency vectors can be used to represent the rate variation encoded in empirical amino acid substitution matrices. This study demonstrates that a rapid and simple method can be developed from the mutation-selection model to predict substitution rates from amino acid data, complementing the standard phylogenetic approach.

## Introduction

Proteins evolve under several genetic, biophysical, and functional fitness constraints(Liberles, et al. 2012). This results in a substantial variation in amino acid substitution patterns at the level of sites in proteins. Amino acid-based phylogenetic models primarily account for this variation by introducing factors that scale the relative substitution rate at sites to the overall mean substitution rate of a multiple sequence alignment (MSA)(Yang 1993; Arenas 2015). Such models have resulted in the improved inference of phylogenic trees(Le and Gascuel 2008), but the individual rate values themselves have also proven to be insightful, in particular when mapped onto three-dimensional structures of proteins to characterize sequence conservation(Pupko, et al. 2002a; Landau, et al. 2005). Site-level data can report on the contribution of sites to the stability, structure, and function of proteins and enables correlation between substitution patterns and molecular properties(Echave, et al. 2016).

Because the evolutionary process proceeds through changes at the nucleotide level, protein models are inherently phenomenological(Liberles, et al. 2012; Jones, et al. 2018). Furthermore, there is significant variability of amino acid propensities at sites in proteins, which cannot be fully accounted for by models that combine site rates with invariable equilibrium frequencies(Echave, et al. 2016). Incompatibilities between the generative process behind protein sequence evolution and inference models used to analyze sequence data can sometimes lead to biased and overconfident predictions(Jones, et al. 2018). By modeling the evolutionary process at the codon level, the mutation-selection model(Halpern and Bruno 1998; Yang and Nielsen 2008; Rodrigue, et al. 2010; Tamuri, et al. 2012) provides an approach to describe site-level heterogeneity with greater realism than protein-level models. Modeling substitution processes at the level of codons and sites requires models with a large number of parameters, which results in computational as well as statistical challenges. Nonetheless, statistical inference frameworks have successfully been implemented to derive site-based parameters from sequence data(Yang and Nielsen 2008; Rodrigue, et al. 2010; Tamuri, et al. 2012). Mutation-selection models have primarily been used to detect signals of adaptive evolution in genes or sites in genes(Tamuri, et al. 2014; Bloom 2017; Rodrigue, et al. 2020).

In the standard approach, the mutation-selection model is applied to nucleotide data. But in many situations, it is preferable to analyze amino acid sequences. By combining the mutation-selection model with a nucleotide mutation model it is possible to predict site-based substitution rates(Halpern and Bruno 1998; Moses, et al. 2003) based on predicted amino acid equilibrium sequences. We show here that this can lead to accurate predictions of site rates from amino acid sequences and improvements relative to the standard phylogenetic approach.

Average substitution rates for amino acids can also be predicted from the mutation-selection model by condensing codon-level rates into amino acid instantaneous rate matrices (Norn, et al. 2020). We demonstrate here that such matrices are very similar to empirical rate matrices like WAG(Whelan and Goldman 2001) and LG(Le and Gascuel 2008). The mutation-selection framework establishes a direct link between amino acid frequency distributions at sites and protein substitution matrices. This connection has been made previously by Pollock and Goldstein, who demonstrated that amino acid substitution rates could be constructed by calculating a covariance matrix between site equilibrium frequency vectors(Goldstein and Pollock 2016). Here we exploit the link between site frequency distributions and substitution matrices to establish which distribution of site frequency vectors is encoded in empirical matrices such as JTT(Jones, et al. 1992), WAG, and LG.

## Results

### Amino acid rates from a mutation-selection model

The mutation-selection model is a codon-based model that describes the effect of protein-level site-specific selection(Bruno 1996). Several simplifying assumptions are made to arrive at the model. Selection pressures at sites are taken to be constant over all lineages, fixation is assumed to be rapid compared to mutation, sites are independent, and the evolutionary process is dominated by negative selection. With these premises the relative instantaneous rate between (*q*_*uv*_) from codon *u* with amino acid *i* to codon *v* with amino acid *j* is modeled as the product of the proposal rate of mutating codon *u* to *v*, *p*_*uv*_, and the fixation probability, *f*_*uv*_, and an arbitrary scaling constant *k*:

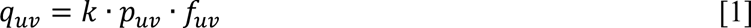

The codon transition probability can be modeled with many approaches, but we assume here that the fitness of a codon is only a function of the amino acid it encodes and that the rate of mutation between codons can be modeled by the nucleotide mutation rate of the nucleotide change separating connected codons (Norn, et al. 2020). Many nucleotide substitution models can be utilized, the results here are obtained with the K80 model with one parameter (*κ*) controlling the relative rate of transitions to transversions (Kimura 1980). In addition, we assign a rate for multi-nucleotide mutations (*ρ*) accounting for insertions, deletions, and tandem mutations that have been shown to play an important role in evolution(Reid and Loeb 1993; Harris and Nielsen 2014). With these parameters, the mutation proposal rates become:

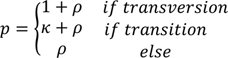

By using the weak mutation model of Golding and Felsenstein (Golding and Felsenstein 1990), the fixation probability *f*_*i,j*_ can be approximated as:

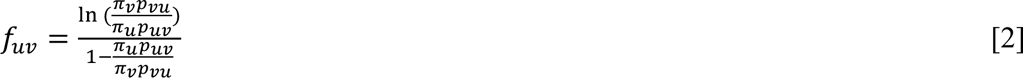

Where π_u/v_ is the equilibrium frequency of codon u/v and p_u/v_ is the probability of mutation between u/v. To estimate the codon equilibrium frequencies π_u/v_ we first estimated the amino acid frequencies from a multiple sequence alignment (MSA) using their maximum likelihood estimates. The frequency of individual codons was then calculating by multiplying the amino acid frequency by the relative frequencies of codons encoding for this amino acid. This relative frequency can be estimated from genomic data. However, these do not correspond to proposal frequencies since they are determined from sequences after selection. Instead, we calculated the relative codon frequencies by solving for the codon equilibrium frequencies in an instantaneous rate matrix where each codon is assigned equal fitness and renormalize the frequencies coding for the same amino acids to sum to 1.

The site-specific rate substitution rate *μ*^*L*^ (the total flux) is:

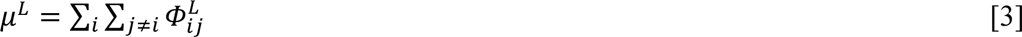

Where *i* and *j* are amino acid types and where 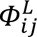 is the flux between i and j which can be found by summing the flux between all connected codons u and v that encode for amino acid i and j:

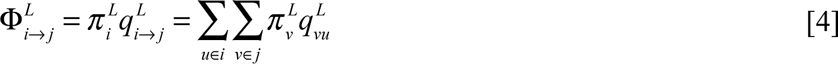

We scale the site-specific rate to correspond to an average of one neutral substitution by per time-unit(Bruno 1996; Spielman and Wilke 2015).

### Maximum-likelihood estimation of model parameters

There are two nuisance parameters in the model, *κ* and *ρ*. Following the approach in Norn et al.(Norn, et al. 2020) we first construct a codon-level instantaneous rate matrix and aggregate it into a protein-level protein rate matrix. To find optimal values of *κ* and *ρ* we find the combination of parameters that maximizes the likelihood in phylogenetic tree reconstruction from a set of MSAs using the corresponding protein rate matrix. Substitution matrices were computed from 221 pfam alignments used by Le and Gascuel to infer the LG matrix(Le and Gascuel 2008). The substitution matrix calculated using each set of parameters was then evaluated by inferring trees 59 TreeBase alignments, also taken from the study by Le and Gascuel(Le and Gascuel 2008). The sum log-likelihoods were used to identify the optimal parameters. A total of 42 parameter combinations were evaluated in a grid search. *κ*=1.4 and *ρ*=0.2 were identified as the optimal solution. The likelihood surface is very flat with respect to the variation of *κ*, whereas a clear optimum was found for *ρ* (**Supplementary Figure 1**). The resulting protein rate matrix can be compared to the empirical rates matrices JTT, WAG, and LG. In **Figure 1**, the amino acid substitution rate values of the optimal rate matrix (referred to as *MutSel*, with *κ*=1.4 and *ρ*=0.2) are compared to the values of JTT, WAG, and LG on the logarithmic scale. There is a strong correlation between the values in these matrices, with r^2^ values of 0.89, 0.83, and 0.79 for JTT, WAG, and LG, respectively. This demonstrates that the relative average amino acid substitution rates are well predicted by the model (to the extent that empirical matrices correspond to true values). However, there is a non-unit slope in the correlation with the empirical matrices that is caused by a lower variation of amino acid rates in the MutSel model.

**Figure 1:**
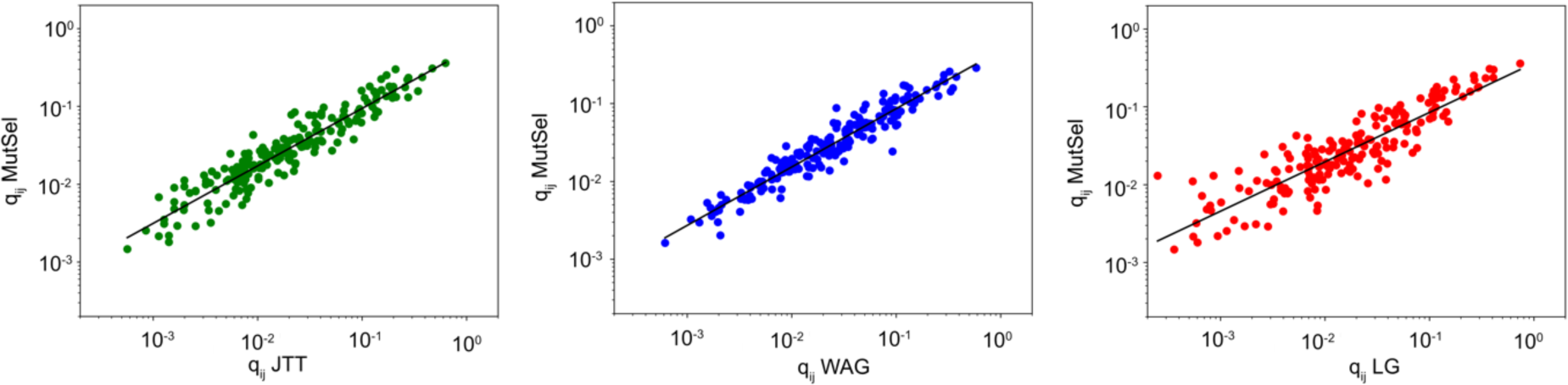
Comparison of MutSel rate matrix with JTT(left), WAG(center) and LG(right). The plot of amino-acid rates (q_ij_) of WAG(left)/LG(right) and the same element in MutSel. Black line, linear fit.

Two aspects could give rise to biased predictions. First, limited sampling of sequences means that the equilibrium frequencies for rare amino acid variants will tend to zero values. This can be combatted by introducing pseudo-counts, which are often used during remote homolog detection(Henikoff and Henikoff 1994). To avoid biasing results towards the empirical matrices we compare to, we tested a simple approach for adding pseudo-counts using a Jeffreys prior for a multinomial distribution. This, however, resulted in a significantly reduced performance. The second aspect is that MSAs do not correspond to random samples of sequences. This can be addressed with the reweighting of sequences before the calculation of equilibrium frequencies. We tested a scheme that weight sequences based on a phylogenetic tree(Stone and Sidow 2007). This resulted in a small, but consistent, reduction in performance.

### Simulation of protein sequence data

Having established values for the nuisance parameters of the model we now go on to evaluate the capability of the mutation-selection model in estimating site-rates. The dataset used in the validation was the result of evolutionary dynamics simulations. A benefit of this simulation strategy is that the ground truth is known in this scenario. For four proteins (Barnase, Ribonuclease T1, Lysozyme, and protein inhibitor CI2) we simulated evolutionary trajectories according to phylogenetic trees inferred from natural MSAs for these systems (data taken from Norn et. al.(Norn, et al. 2024)). In the simulations, a recursive algorithm was used to “crawl” the tree and at each new node an evolutionary dynamics trajectory is launched with a branch length taken from the empirical tree. RosettaEvolve(Norn and Andre 2023), part of Rosetta macromolecular modeling package(Leman, et al. 2020), was used for protein structure-based evolutionary dynamics simulations. RosettaEvolve is similar to other structure-based evolutionary dynamics methods presented in the literature(Goldstein 2011; Pollock, et al. 2012; Shah, et al. 2015; Jiang, et al. 2018). Mutations are proposed at the nucleotide level and correspond to either single-base pair mutations or whole-codon codon changes. The probability of fixating a mutation is estimated using Kimura’s expression for fixation probability(Kimura 1962). The simulations are carried out with the assumption that neutral evolution is primarily driven by thermodynamic stability, with a fitness function that equals the fraction of folded protein(Williams, et al. 2006). The effect of mutations on stability is calculated using structure-based ΔΔG calculations(Park, et al. 2016). The sequence diversity generated in these simulations is depending on the assumed stability of the protein. The stability was selected in these simulations to result in sequence entropies of simulated alignments that is similar to the values in the empirical MSA. The leaf-nodes were collected to create MSAs from which site-rates were predicted. Because the full evolutionary trajectories during the tree crawling is recorded the true substitution rate is known for each site in a protein.

The comparison of substitution rates estimated from the mutation-selection model and substitutions counted per site counted from the trajectories are shown in the upper row in **Figure 2** for the four simulated proteins.

**Figure 2:**
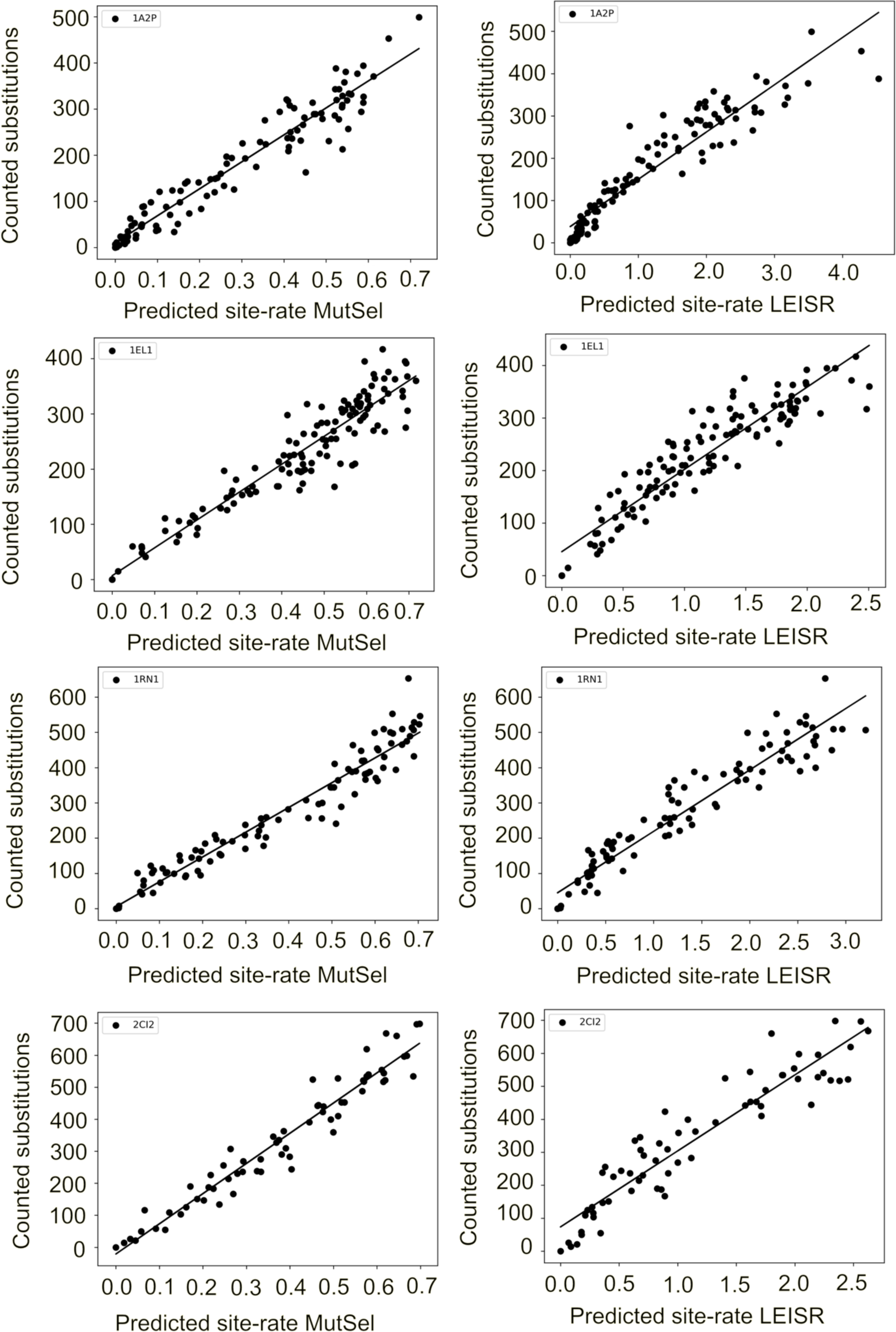
Comparison of rates predicted from sequence and the true substitution count. Left column, comparison of site rates predicted from the mutation-selection model compared with counted substitutions in evolutionary dynamics simulations based on the structure of Barnase (PDBID: 1a2p), Ribonuclease T1 (PDBID: 1rn1), Lysozyme (PDBID: 1el1) and CI2 (PDBID: 2CI2). Black circles, individual site-rates. Black line, linear fit. Right column, a comparison of site rates predicted using phylogenetic inference through LEISR(Spielman and Pond 2018) to counted substitutions for 1a2p, 1rn1, 1el1, and 2ci2. Black circles, individual site-rates. Black line, linear fit.

The rates predicted from the mutation-selection model have a strong linear correlation with the counted substitutions from the evolutionary dynamics simulations. r^2^ values range from 0.90-0.94 (Barnase: 0.93, Ribonuclease T1: 0.94, Lysozyme: 0.90, CI2: 0.94). We compared these results with site rates inferred using Maximum likelihood calculations using LEISR(Spielman and Kosakovsky Pond 2018), which builds upon the methodology of the pioneering rate4site(Pupko, et al. 2002b), but has improved support for larger MSAs. LEISR also accurately predicts site rates (bottom row **Figure 2**), with r^2^ values ranging from 0.87-0.92 (Barnase: 0.88, Ribonuclease T1: 0.92, Lysozyme: 0.87, CI2: 0.88). In contrast to the mutation-selection model the correlation counted substitutions deviates slightly from linear correlation.

The predictions from the mutation-selection model and LEISR were made with relatively large MSAs (Barnase: 237 sequences, Ribonuclease T1: 306, Lysozyme: 537, CI2: 529). Because the estimation of the equilibrium frequencies should become more uncertain as the number of sequences decreases, we expect the performance of the site-rate prediction to deteriorate. To investigate the correlation of accuracy of site-rate predictions as a function of the depth of the sequence alignment we sampled smaller subsets of sequences and predicted site-rates from these. **Figure 3** shows how the r^2^ between predicted rates and counted substitutions varies with the depth of the sequence alignment.

**Figure 3:**
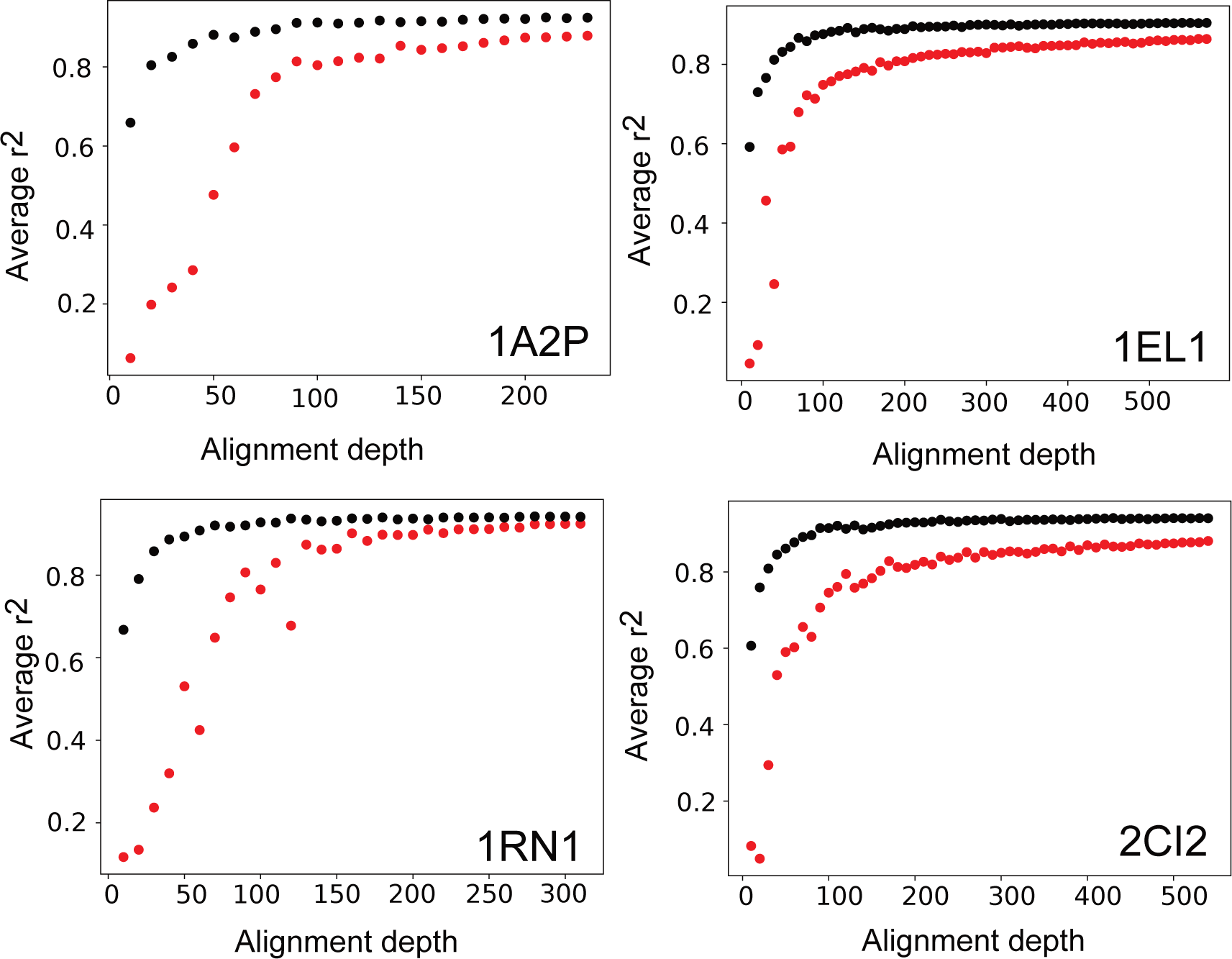
Effect of depth of MSA on site-rate prediction. Average Pearson r^2^ between predicted rates from the mutation-selection model (black) and LEISR (red) as a function of the number of sequences in the alignment. Each point is an average of 5 random sets of sequences.

The results show that 50 or more sequences are required to reach the highest correlation with the empirical substitution counts for the mutation-selection model. Nonetheless, with 30 sequences the r^2^ values are still around 0.8 demonstrating that accurate predictions can be made with relatively few sequences. In these examples, LEISR requires more than 125 sequences to reach similar accuracies and has consistently lower correlation across all sequence depths.

### Comparison with rates inferred by maximum likelihood

Proteins evolve by an evolutionary mechanism that is considerably more complex than what is encoded in simulated data. To investigate how the mutation-selection model predicted rates from a set of 213 protein sequence alignments presented by Echave et. al. (Echave, et al. 2015). This study included site rates estimated using the empirical Bayes rate4site method, using two different substitution matrices, JC and JTT. In **Figure 4** the correlation between rates estimated from the mutation-selection model and the rates4sites are shown.

**Figure 4:**
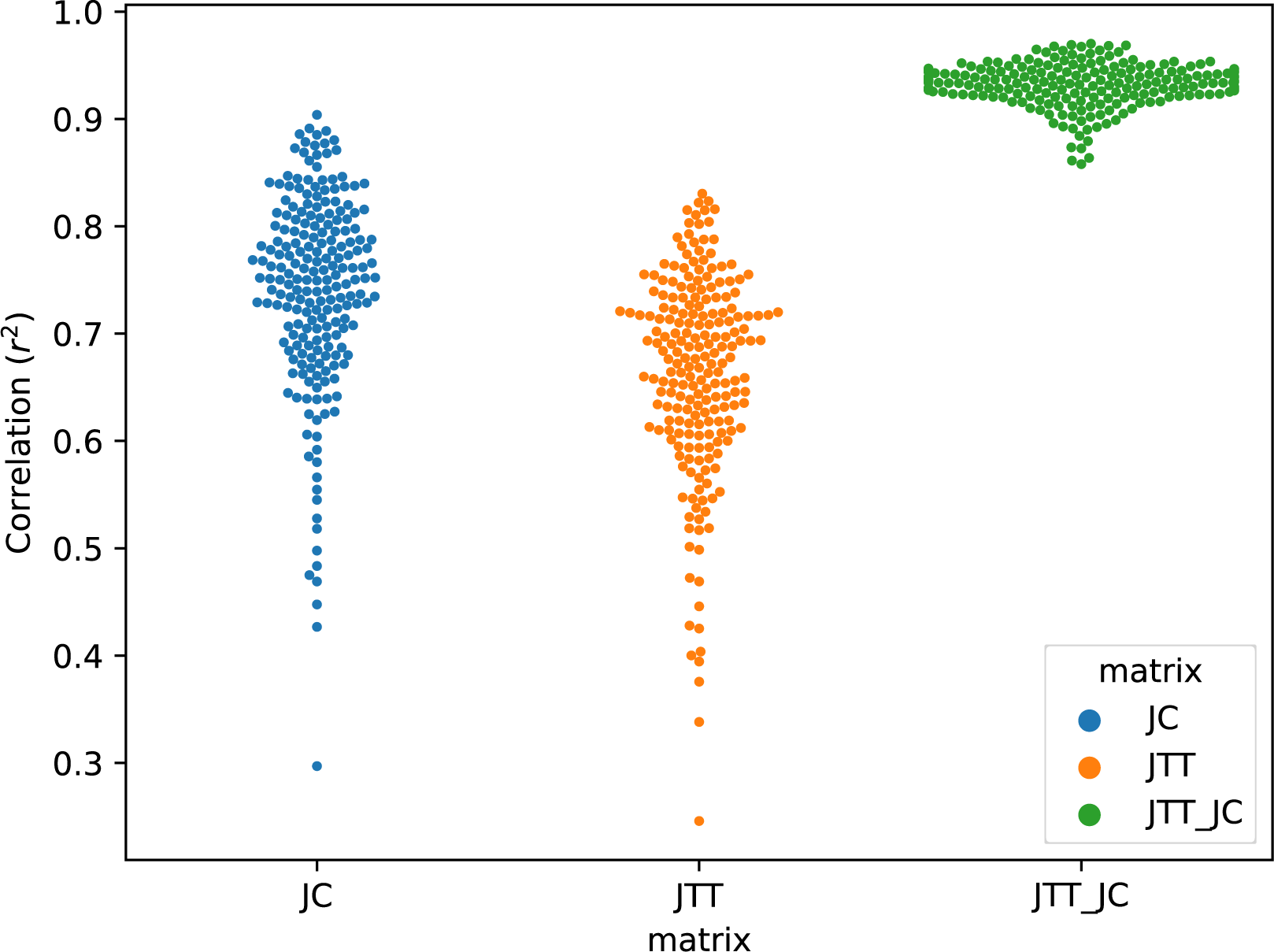
Correlation between rates from mutation-selection model and rates for site. Site-rates estimated from 213 MSAs using the mutation-selection model and rate4sites with two different substitution matrices (JC and JTT, blue and orange respectively) and compared with r^2^ correlation in swarmplot. Correlation between rates estimated using JC and JTT are shown in green.

The average r^2^ correlation is 0.74 to rates inferred with the JC matrix and 0.66 to JTT. The amino-acid JC model(Mayrose, et al. 2004) assigns an equal probability for all amino acid mutations, which gives higher correlations with the mutation-selection. The choice of substitution matrix has an impact on inferred rates (average r^2^ 0.93), but this variation is smaller than the variation between mutation-selection model and maximum likelihood. Nonetheless, the mutation-selection model gives similar rates to empirical Bayes.

### Deconstructing empirical substitution matrices into the distribution of equilibrium frequency vectors

The mutation-selection model couples the distributions of equilibrium frequencies to the amino acid substitution rates in instantaneous rate matrices. The site frequency vectors used to construct an amino acid substitution matrix constitute an ensemble of site probability vectors. Such ensembles will be unique for each empirical substitution matrix. In this section, we derive such probability vectors for JTT, WAG, and LG by fitting a distribution that maximizes the correlation between the mutation-selection matrix and the empirical matrix. Such deconvolution can give insight into what the implicit assumptions about site rates have gone into empirical rate matrices used in phylogenetic analysis.

Previous studies have demonstrated that the amino-acid frequency distribution at sites in proteins is well approximated by the following expression(Porto, et al. 2005; Johnson and Wilke 2020) if the frequencies (π) are ordered from large to small values (π_1_>π_2_…>π_20_):

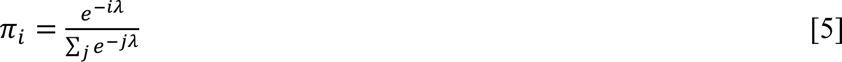

where *i*=1..20 and λ controls the shape of the distribution of ρε-values. Large values of λ results in frequency profiles dominated by a few amino acids. We used the method presented in Johnson and Wilke(Johnson and Wilke 2020) and determined a histogram of λ-values in the MSAs used to fit the LG matrix. The λ-distribution is well-described by a gamma function (**Supplementary figure 2**). We sampled 1000 values from this gamma λ-distribution to create an ensemble of frequency vectors. Equation 5 does not provide a unique mapping to amino acid identities. We used a sampling approach to assign identities of amino acids to these frequency vectors so that they approximately resulted in average frequencies of amino acids that matched the equilibrium frequency distribution in LG. From this ensemble of 1000 frequency vectors, a subset of 100 vectors were picked out that were used to calculate an instantaneous rate matrix through the mutation-selection framework. The ensemble was selected to maximize the correlation with the empirical matrices JTT, WAG, and LG. Optimization was carried out by simulated annealing. The result of this procedure is an ensemble of frequency distributions that maximizes the correlation to empirical matrices and the result is shown in **Figure 5**.

**Figure 5:**
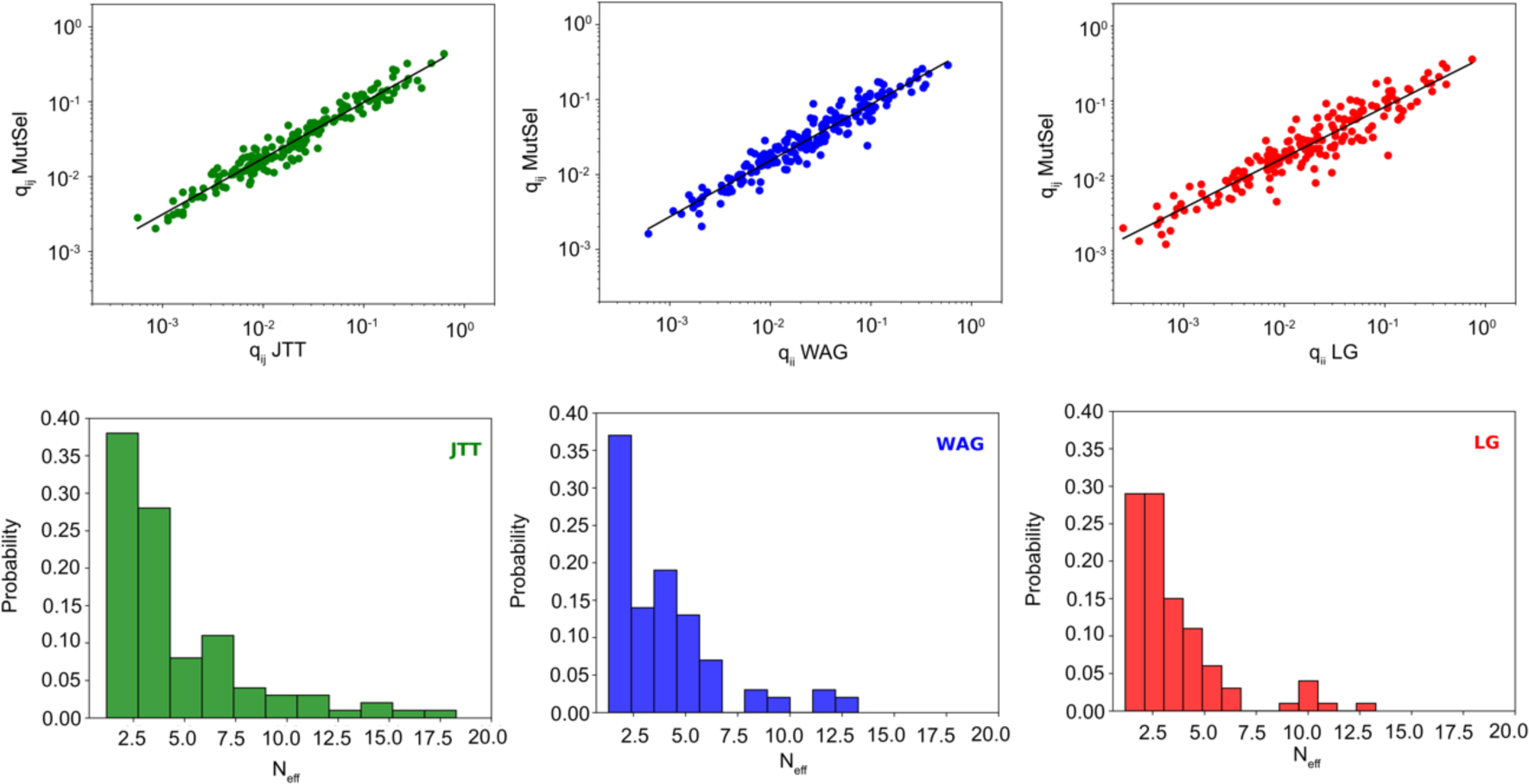
Fit of amino acid frequency distributions to empirical matrices. Upper row, plot of amino-acid rates (q_ij_) of JTT(left)/WAG(center)/LG(right) and the same element in fitted mutation-selection model. Black line, linear fit. Lower row, histogram of the effective number of amino acids (N_eff_) in fit to JTT, WAG, and LG.

The fitted matrices are highly correlated to their empirical counterparts with r^2^ values of 0.95, 0.92, and 0.87 for JTT, WAG, and LG, respectively. To characterize the fitted frequency distributions, the effective number of amino acids (n_eff_) was calculated for each vector in the ensemble. This is a convenient metric to characterize variability at sites in a protein. n_eff_ values range between 1 and 20, where 1 means that only one type of amino acid is found at a site and 20 means that all 20 occur with equal probability. The distribution of effective amino numbers for each matrix is presented in **Figure 5**. JTT is substantially different from WAG and LG, with a broader distribution and more vectors corresponding to highly variable “sites” (higher n_eff_ values). The average n_eff_ for JTT is 4.6. The distribution of WAG and LG have more in common, although WAG has a higher average n_eff_ than LG (4.0 vs 3.4). This primarily stems from the higher frequency of very invariable “sites” for WAG (low n_eff_ values).

## Discussion

The mutation-selection model provides a powerful framework for describing how the action of mutation and selection combine to generate site variability in protein sequences. In this model, site variability emerges as a consequence of the relative time-averaged fitness of codons and individual amino acids. Phylogenetic models used in tree inference from protein sequences typically assume that all sites evolve with the same equilibrium frequencies and site variability is introduced by a phenomenological rate scale factor applied to each site. This restriction can be loosened by allowing the equilibrium frequency distribution to vary at sites (Quang le, et al. 2008). However, the large number of parameters of such models coupled to their computational complexity limits their utility. The performance of variable site frequency models in site-rate predictions has also not been benchmarked.

We focus here on the prediction of substitution rates at individual sites of proteins from amino acid sequence data and show that they can be calculated using the mutation-selection framework with just two nuisance parameters (*κ* and *ρ*). In practice, the model is insensitive to the value of *κ* leaving a single modeling parameter. The main complication is the estimation of equilibrium frequency values from MSAs. The amino acid distributions of leaf sequences are not necessarily good estimators of equilibrium frequencies. For sequences with limited divergence, this may become an issue, limiting the approach presented here. There is also an issue with uncertainties in equilibrium frequency distributions due to limited statistical sampling. However, based on experiments using a simulated data set, we showed that accurate predictions of site rates can be predicted with as little as 20 sequences for two of the systems studied here. This is surprising but can be explained because most sites are highly conserved so equilibrium frequencies can be estimated with relatively few sequences. Also, since site rates are an aggregate of rates across many codons, errors in individual amino acid frequency values may be washed out in the model. Comparison with rate-scale factors predicted from phylogenetic inference demonstrates some benefits of the mutation-selection model. Prediction accuracy is better for all simulated systems and across all alignment depths and computations can be carried out much more rapidly. Nonetheless, with enough sequences, the maximum likelihood estimated rate factors are also accurate. Since the underlying data is based on nucleotide mutation trajectories, it is encouraging to see that both approaches result in good predictions.

Because benchmarks with known rates are difficult to construct from empirical data, we have used a simulation approach in this study. The evolutionary dynamics simulation generates sequences that result from time- as well as context-dependent fluctuations of fitness. By simulating evolutionary dynamics using a phylogenetic tree, sequences maintain similar phylogenetic relationships as those found in natural sequences. The approach thus provides a more realistic model of protein evolution than nucleotide or amino acid-based simulators. However, one caveat is that the mutation model underlying the evolutionary dynamics simulations shares similarities to the one used in the mutation-selection model, which could give the mutation-selection model an advantage in the comparisons. In particular, the proposal rates in the evolutionary dynamics simulations are also governed by a single nucleotide mutation rate *κ* and a multi-nucleotide rate *ρ*. However, the parameter values used in the evolutionary dynamics simulation were different from the values used to calculate the site rates and we show that the value of *κ* does not influence the rates much. One factor not included in the evolutionary dynamics simulation is selection at the codon level, which we know occurs in natural sequence evolution. Such codon-fitness models could be introduced into the mutation-selection framework, further improvement could also be made by using a more complex, but parameter-rich, model for nucleotide mutations. However, it is not clear that this would improve the result given the fact that the input data is amino acid sequences that do not inform on codon usage. With more parameters in the model, using a continuous method for maximum likelihood optimization of parameters would be beneficial(Holder, et al. 2008). The accuracy of site-rate predictions has been previously characterized by Scheffler et al. (Scheffler, et al. 2014). They found that rates predicted from phylogenetic inference were linearly correlated with experimental values and that inference showed good convergence. However, one caveat with their study is that rates were simulated with the same gamma-distributed rate factors used in the inference model.

The matrix fitting result demonstrates that mutation-selection model can be parametrized to give rise to matrices that are very similar to the empirical rate matrices, with r^2^ values going up to 0.95 for JTT. This means that we can think of substitution matrices as being the result of an averaging of an ensemble of site-specific rate matrices, each parametrized by a single equilibrium frequency vector. Nonetheless, there appears to be an upper limit to the quality of the fit because independent fitting experiments converge to similar correlation values. The more complex the matrix (JTT<WAG<LG), the lower the correlation (same as the pattern for the matrices calculated from natural sequences in **Figure 1**). LG, for example, differs from WAG by being inferred by introducing site-rate scale factors in the maximum likelihood estimation. Because of the vast number of parameters in the WAG and LG fitting and statistical uncertainty in inference, it is not expected that mutation-selection model could explain the full variation in the empirical matrices.

The fitted distributions of equilibrium frequencies can be used to understand some of the implicit assumptions encoded in empirical matrices. JTT is constructed by the traditional method of counting amino acid substitutions in MSAs(Jones, et al. 1992). JTT is consistent with a broad ensemble of highly variable site frequency vectors. The introduction of site-specific rate factors in LG compared to LG results in a lower frequency of highly conserved sites. By varying the ensemble of frequency vectors, it is possible to create a vast array of substitution matrices with different properties.

In this study, we have demonstrated that substitution rates at sites in proteins can be calculated by combining the mutation-selection model with a nucleotide mutation model and equilibrium frequency predictions from protein sequence alignment. This was used by Halpern and Bruno(Halpern and Bruno 1998) in the original mutation-selection study to characterize branch lengths but has not been generally applied for site-rate predictions from amino acid data. The approach enables fast and accurate site-rate predictions even with limited sequence data and provides a link between the frequency distribution at sites in proteins and amino acid substitution rates encoded in amino acid substitution matrices.

## Materials and Methods

### Rate matrix calculations

The 3912 pfam alignments employed by Le and Gascuel to train the LG matrix(Le and Gascuel 2008) were used as the basis for calculating protein rate matrices. We selected a subset of alignments with 30 or more sequences, leaving 221 MSAs. Amino acid frequencies at each site were estimated by counting the occurrences in each column of the MSA, not counting gaps. Sites with more than 50% gaps were discarded, although this filter had no substantial impact on the result. To account for deficiencies in statistical sampling pseudo-counts are often employed. When pseudo-counts were tested, we used a Jeffreys prior and added a count 0.5 to each amino acid. To account for biased sampling of amino acid sequences we weighted the sequence based on a phylogenetic tree. For each of the 221 alignments we use FastTree(Price, et al. 2010) to calculate phylogenetic trees. BranchManager(Stone and Sidow 2007) was then utilized to create weights for each sequence based on the tree. A codon-based instantaneous rate matrix was then constructed based on equation 1. By assigning equal fitness to codons in the model we can solve for the equilibrium frequency distribution and normalize the frequencies so that codons that code for the same amino acid sum to 1. Codon frequencies were then calculated by multiplying the relative codon frequencies with the amino acid frequency estimated from the MSA. Following the aggregation procedure in Yang et al.(Yang, et al. 1998) and Norn et al.(Norn, et al. 2020) the codon-level instantaneous rate matrix can be by condensed into a protein-level matrix. The instantaneous rate (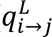) between two amino acids (*i*, *j*) with codons *u*∊*i* and *u*∊*i* can be calculated as:

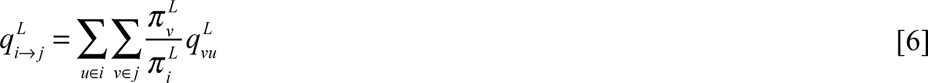

Where *π* is an equilibrium frequency. By averaging the flux at many sites in many protein MSAs we can calculate the mean instantaneous rate for amino acid *i* to *j* as:

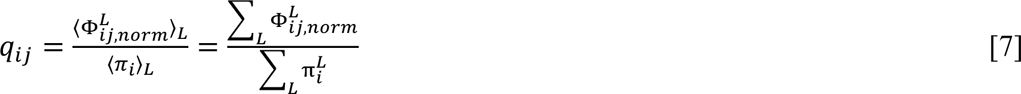

Before averaging, the site rates were normalized to correspond to one neutral substitution by per time-unit(Bruno 1996; Spielman and Wilke 2015). Rate calculations take on the the order of 10s.

#### Maximum Likelihood estimation of model parameters

A grid-based Maximum Likelihood search was carried out for the parameters, (*κ* ∈ [1.4-2.6] and *ρ* ∈ [ 0-1.0]. For each combination *κ* and *ρ* values we generated a protein rate matrix. The rate matrix was used for the inference of phylogenetic trees, using the 59 TreeBase test alignments utilized by Le and Gascuel in their study on the LG matrix(Le and Gascuel 2008). IQTREE(Minh, et al. 2020) with MATRIX+FO+G4 (Where MATRIX is the matrix mutation selection matrix) was used for tree inference. The likelihoods were summed up for all 59 alignments to identify the maximum likelihood solution.

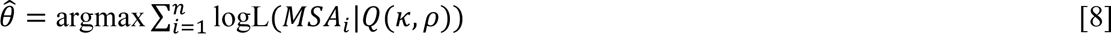

#### Evolutionary dynamics simulations

Evolutionary dynamics were simulated with RosettaEvolve, which is part of the Rosetta macromolecular modeling package(Leman, et al. 2020). The data in this study was taken from Norn et. al.(Norn, et al. 2024), which together with the study presenting RosettaEvolve(Norn and Andre 2023) gives a complete methodological description. Phylogenetic trees for simulations were generated using RaXML(Stamatakis 2014) based on natural sequence alignments of the four proteins simulated in the study. Tree-crawling started from the center of the tree by simulating each individual branch of the tree in a recursive fashion. The leaf nodes were then used for site-rate predictions.

### Site-rate fitting of empirical matrices

The 3912 sequences in the pfam sequences used to fit the LG matrix was analyzed to extract a histogram over λ-values (equation 5) with scripts and methods presented by Johnson and Wilke(Johnson and Wilke 2020). A gamma distribution was fitted to the λ-values, results shown in **Supplementary Figure 2**, using the *fitdistr* package in R(Team 2020). The resulting gamma distribution was used to sample 1000 λ-values. Equation 5 provides a list of sorted frequency values from large to small. To assign identities to a frequency vector a simple approximate sampling approach was used. For the first largest frequency value in the vector amino acids are sampled according to the equilibrium frequencies of LG. Then the second largest element is sampled, with frequency according to the equilibrium frequencies of LG excluding the already selected amino acid. This is iterated until the 20^th^ amino acid is assigned. The resulting vectors have mean probabilities that differs slightly from LG. An ensemble of these can selected to perfectly converge to the LG equilibrium frequencies. Since the ensemble will anyway be fitted in the next step, we skipped this step. Simulated annealing (https://github.com/perrygeo/simanneal) was then used select a 100 site-frequency vectors out of the ensemble of 1000 vectors to optimize the Pearson-correlation between a matrix calculated from the ensemble and a reference matrix (JTT,WAG or LG).

Effective number of amino acids was calculated as:

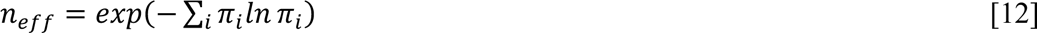

### Data availability

*The data underlying this article are available in*https://github.com/Andre-lab/mutsel-rate-manuscript.git. or *will be shared on reasonable request to the corresponding author*.

