## Supplemental materials for "Accurate prediction of site- and amino-acid substitution rates with a mutation-selection model"

**Ingemar André[1]**

*[1]Biochemistry and Structural Biology, Lund University, PO BOX 124, Lund, Sweden*

**

### Supplementary Information

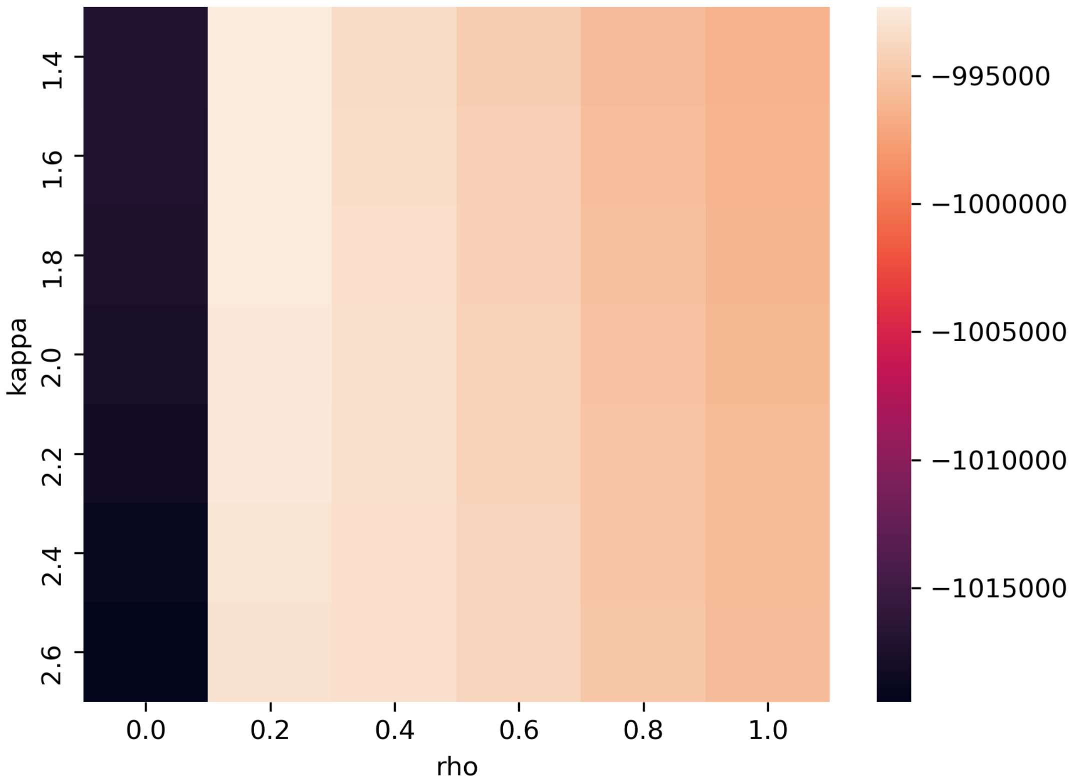


***Supplementary figure 1:*** *Maximum likelihood estimation of κ and ρ. Heatmap of summed likelihood for 59 alignments as function of κ and ρ.*

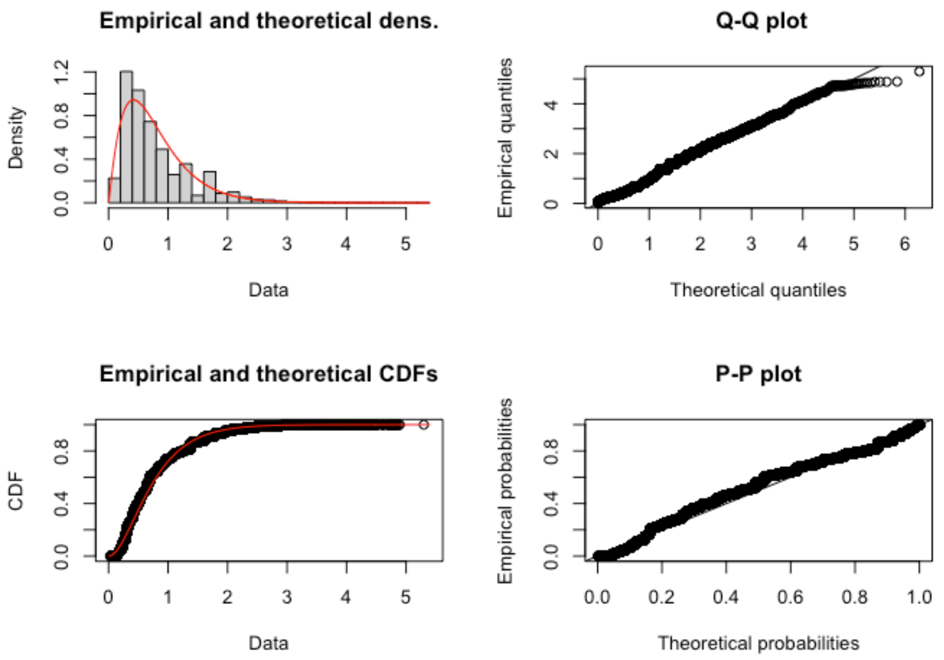


***Supplementary figure 2:*** *Fit of λ-distribution to a gamma function. Fitting result from fitdistr package in R(Team 2020) of the empirical histogram of λ-values from the multiple sequence alignments used to fit the LG substitution matrix.*
